# Dynamics of interaction networks and species’ contributions to community-scale flexibility

**DOI:** 10.1101/2022.12.11.519979

**Authors:** Hirokazu Toju, Sayaka S. Suzuki, Yuki G. Baba

## Abstract

Architecture of species interaction networks is a key factor determining stability of ecological communities. However, the fact that ecological network architecture can change through time is often overlooked in discussions on community-level processes despite its theoretical importance. By compiling a time-series community dataset involving 50 spider species and 974 Hexapoda prey species/strains, we quantified the extent to which architecture of predator–prey interaction networks can shift across time points. We then developed a framework for finding species that could promote flexibility of interaction network architecture. Those “network coordinator” species are expected to promote persistence of species-rich ecological communities by buffering perturbations to communities. Although spiders are often considered as generalist predators, contributions to network flexibility varied greatly among species. We also found that detritivorous prey species can be cores of interaction rewiring, dynamically interlinking below-ground and above-ground community dynamics. Analyses of network coordinators will add a new dimension to our understanding of species coexistence mechanisms and provide platforms for systematically prioritizing species in terms of their potential contributions in ecosystem conservation and restoration.

**Significance Statement:** Like networks of human relations, webs of interactions between species are dynamically restructured through time. By compiling time-series time-series dataset including > 1,000 species/strains, we quantified the magnitude of ecological network dynamics in the wild. The analytical framework developed in this study highlighted “network coordinator” species, which are keys to conserve and restore endangered ecosystems.

## Introduction

In nature, numerous species form entangled webs of interactions (1), collectively driving community-scale dynamics (2–5). Since May’s seminal work on relationship between community complexity and stability (6), potential mechanisms by which species-rich communities are maintained have attracted scientists. Mathematical models with simple assumptions basically predict that species-rich ecological communities are inherently unstable (i.e., likely to collapse after perturbation) (6, 7). However, it has been proposed that introducing key features of real ecological communities can reorganize our knowledge of how species-rich communities are maintained (8–12).

One of the key properties of ecological communities is architecture of interaction networks (13–16). In classic mathematical models of community dynamics, interactions have been assumed in randomly selected pairs of species in a community (6, 7). Meanwhile, studies on empirical datasets of species interactions have revealed that community-scale organization of interactions are never random (13, 15, 17, 18). Theoretical studies assuming non-random networks have then predicted that specific types of network architecture, such as nested or compartmentalized architecture, can increase/decrease community stability (11, 19, 20). Although application of network science has significantly promoted theories on species coexistence, majority of studies have still relied on unrealistic assumptions of ecological interaction networks. Specifically, fixed architecture of interaction networks has been often assumed in theoretical and empirical investigations on ecological networks.

The concept that ecological network architecture can dynamically change in nature is central to our understanding of species coexistence in real ecological communities (21–25). If a food web has a fixed (rigid) network structure, extinction or population decline of a predator species may trigger the burst of the population size of some prey species, resulting in extinctions of competing prey species and subsequent cascade extinctions through the interaction network (26–28). In contrast, if species interactions in a food web is flexible (29–31), ecological effects of the extinction of a predator species can be buffered by prey range shifts or functional responses of other predator species (12). Thus, flexibility of interaction networks is considered as the key factor determining community stability (12, 31, 32). Nonetheless, due to difficulty in obtaining time-series datasets of species interactions, few studies have revealed how network architecture can shift thorough time in species-rich communities.

We here examine whether species-rich predator–prey communities can show drastic network architectural dynamics in nature. By compiling an empirical dataset involving 50 spider (predator) species and 974 prey species/strains (33), we quantitatively evaluate shifts of network architecture across eight months. We then decompose the network architectural changes into two components, namely, network shifts due to interaction rewiring and those due to species turnover. Based on the analysis of network dynamics, we develop a framework for evaluating the extent to which respective species in a community contribute to the flexibility of network architecture. In the framework, species with high potential impacts on network flexibility is of particular interest because those species could buffer environmental perturbations to communities. Overall, we propose that insights into such “network coordinators” will reorganize our understanding of mechanisms by which keystone species govern coexistence in species-rich communities.

## Results

### Analyses of network dynamics

Changes in network architecture can be evaluated based on *β*-diversity metrices, which are frequently used in measuring species compositional dissimilarity between local communities (25, 34, 35). When network data matrices of multiple time points or multiple local communities are available, we can calculate not only dissimilarity (*β*-diversity) in species (vertex) compositions (*β*_S_) but also that in interaction (edge) compositions (*β*_INT_) for each pair of community matrices (25). Dissimilarity in interactions can be then decomposed into two components, specifically, network architectural dissimilarity due to interaction rewiring (*β*_RW_) and that due to species turnover (*β*_ST_) as detailed in previous studies (25, 34, 35) (see Material and Methods for details; Fig. 1A).

**Figure 1.**
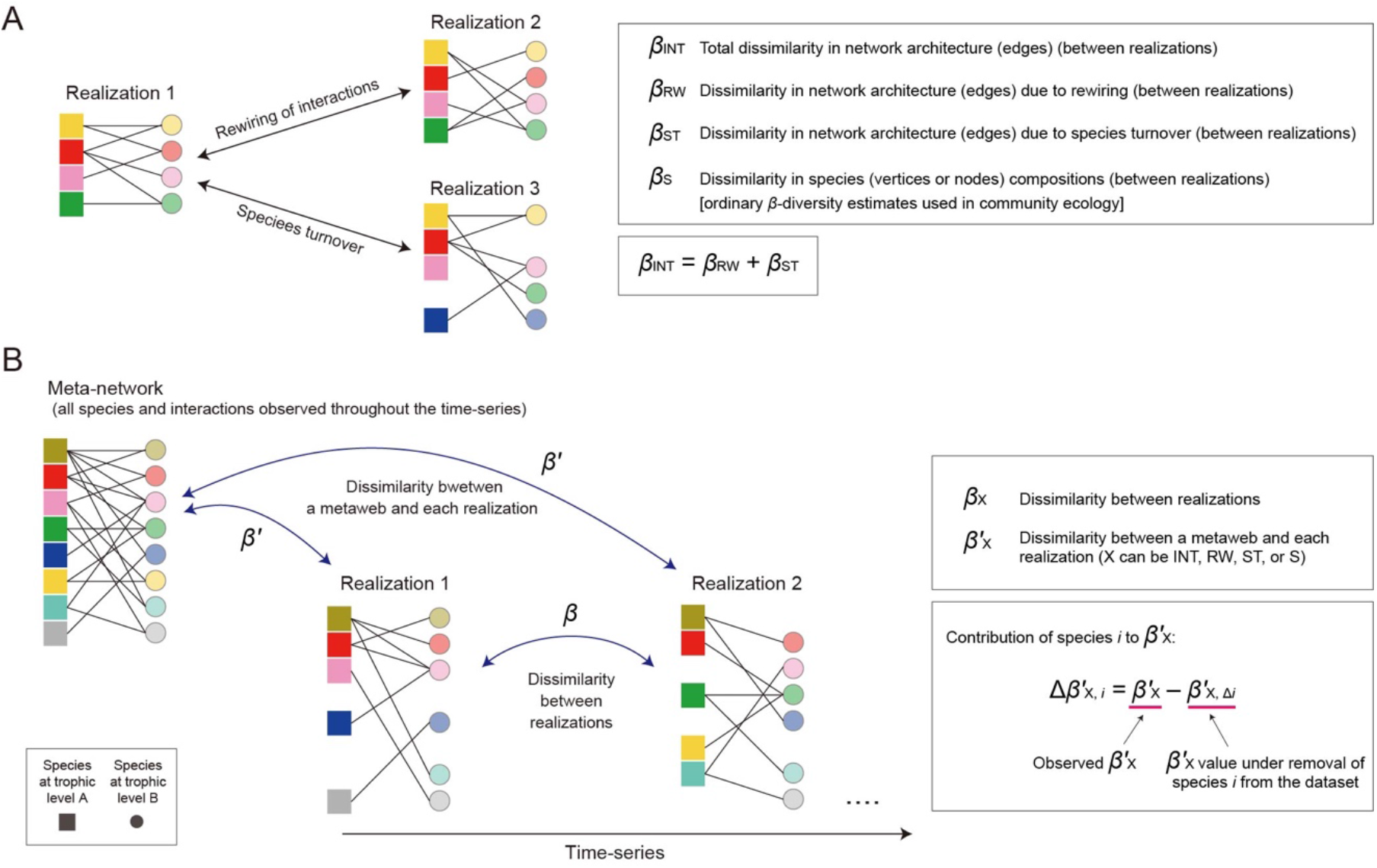
Evaluating dissimilarity in the architecture of species interaction networks. (A) Network architectural dynamics. Both species compositions (network vertices) and interactions (network edges) can vary among communities realized at specific time or space (“realizations”). (B) Meta-networks and realizations. Compiling the information of all the realizations yields the data matrix of the “meta-network” including all nodes and interactions observed through a defined period of time or across metacommunities. In this study, dissimilarity (*β*-diversity) between realizations is designated as *β*, while dissimilarity between a realization and the meta-network is described as *β*^′^. Subscripts of *β* and *β*^′^ represent targets of dissimilarity analyses: “INT”, total dissimilarity in network architecture; “RW”, dissimilarity in network architecture due to rewiring of species interactions; “ST”, dissimilarity in network architecture due to species turnover; “S”, dissimilarity in species compositions are represented. Contributions of each species to respective *β*-diversity components are evaluated as shown in the box.

In addition to comparison between different time points or local sites, comparison of network architecture can be performed between each community matrix representing species interactions realized at specific time or space (hereafter, realizations) and the “meta-network” matrix consisting of all the species interactions observed across the realizations (Fig. 1B). Specifically, for each pair of a realization and the meta-network, total dissimilarity in network architecture 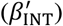, dissimilarity in network architecture due to rewiring 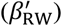, and dissimilarity in network architecture due to species turnover 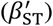 can be calculated (25).

Based on the platform, we here propose a framework for evaluating potential contributions of each species to the flexibility of ecological network architecture. The extent to which dissimilarity in network architecture between a realization and the meta-network can change due to interaction rewiring effects of species *i* are quantified as:

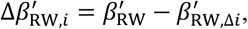

where 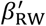 was the original value of dissimilarity in network architecture due to rewiring as defined above, and 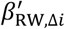denoted the simulated value of dissimilarity calculated by removing species *i* from the dataset (Fig. 1). By definition, this 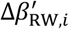index can be calculated for each species in the dataset in each pair of a realization and the meta-network. Therefore, for each species *i*, the maximum value of 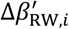across the dataset [i.e., 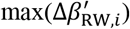] are used as a measure of potential magnitude of contributions to network architectural flexibility. In addition to 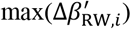, we can calculate 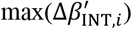, which represented potential magnitude of contributions to total network dissimilarity between realizations and the meta-network (see Materials and Methods for detailes).

### Transitions of network architecture

We first evaluated the extent to which network architecture could change through time by compiling the time-series dataset of predator–prey interactions in a warm-temperate grassland (33). The dataset based on high-throughput DNA metabarcoding included 50 spider (predator) species and 974 prey Hexapoda species/strains in its meta-network (Fig. 2A; Fig. S1), which consisted of eight realizations of predator–prey interaction networks observed from April to November (Fig. 2B). Because the network data included frequency information of each edge (i.e., the number of predator samples from which a focal prey was detected), we used a *β*-diversity metric for quantitative data in the calculations of the indices discussed above. Specifically, Bray-Curtis metric of *β*-diversity was applied after converting frequency information into proportions. Thus, the term “network architecture” represents not only the presence/absence of network edges (i.e., network topology) but also organization of interaction strength (i.e., edge weights) in this study. This assumption of network architectural dynamics is compatible with that of theoretical studies incorporating changes in interaction coefficients (i.e., functional responses) as essential factors determining community stability (12).

**Figure 2.**
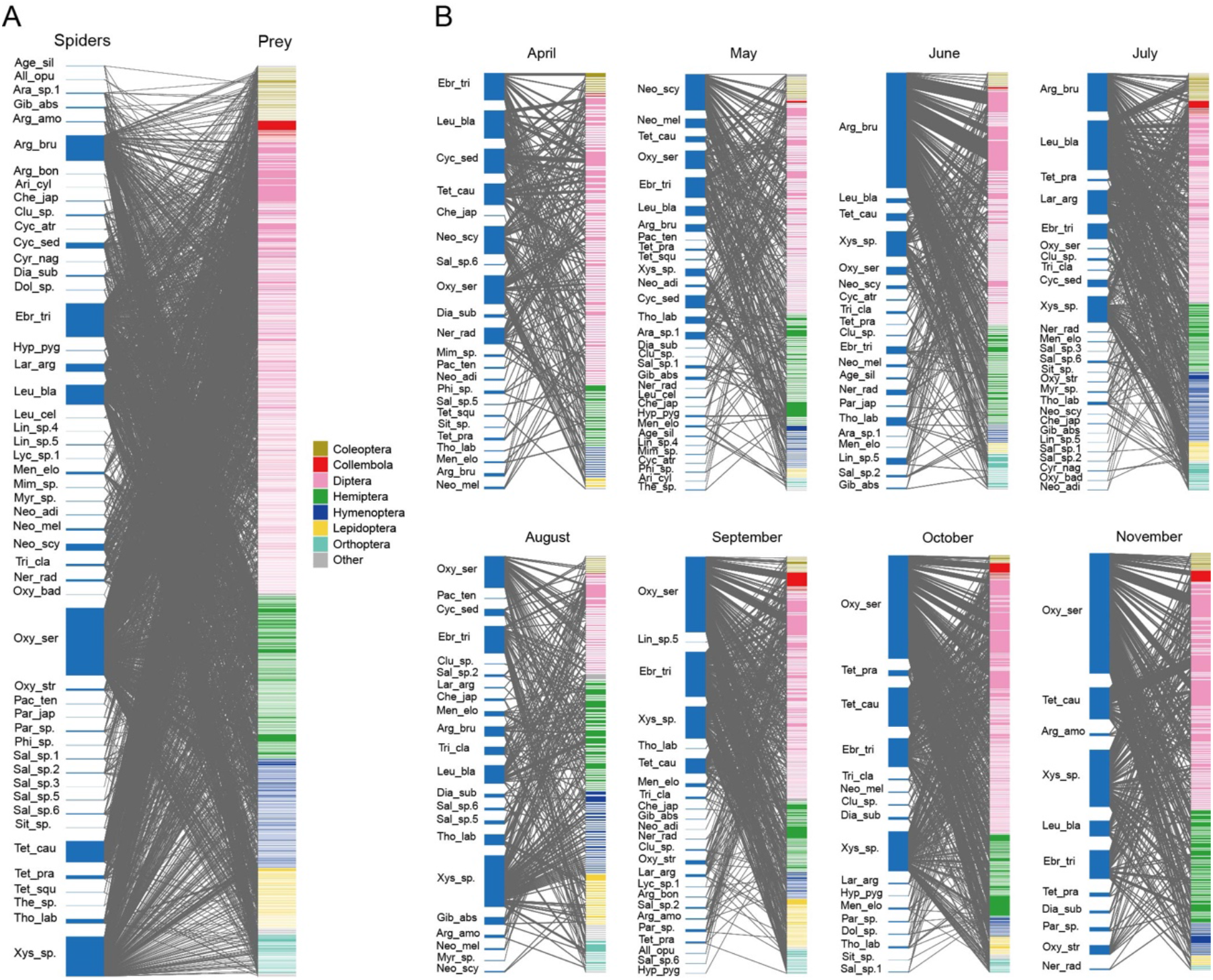
Topology of spider–prey networks. (A) Meta-network including all the spider–prey interactions observed from April to November. Spider species and prey Hexapoda OTUs are shown in the left and right, respectively. See Figure S1 for the abbreviations of spiders and the information of prey OTUs. (B) Network topology of each month. Reproduced from the time-series dataset of Spider–Hexapoda interactions. Reproduced from the data of a previous study (33).

In the spider–prey system, both species (vertex) compositions (*β*_S_) and interaction (edge) compositions (*β*_INT_) continually shifted between consecutive months from April to November (Fig. 3A). In terms of network architectural shifts, not only changes due to species turnover (*β*_ST_) but also changes due to interaction rewiring (*β*_RW_) played major roles. The proportion of interaction rewiring effects to total changes in network architecture (*β*_RW_/*β*_INT_) varied from 28.6 % to 61.0 %, showing the lowest and highest values in April and October, respectively (Fig. 3B).

**Figure 3.**
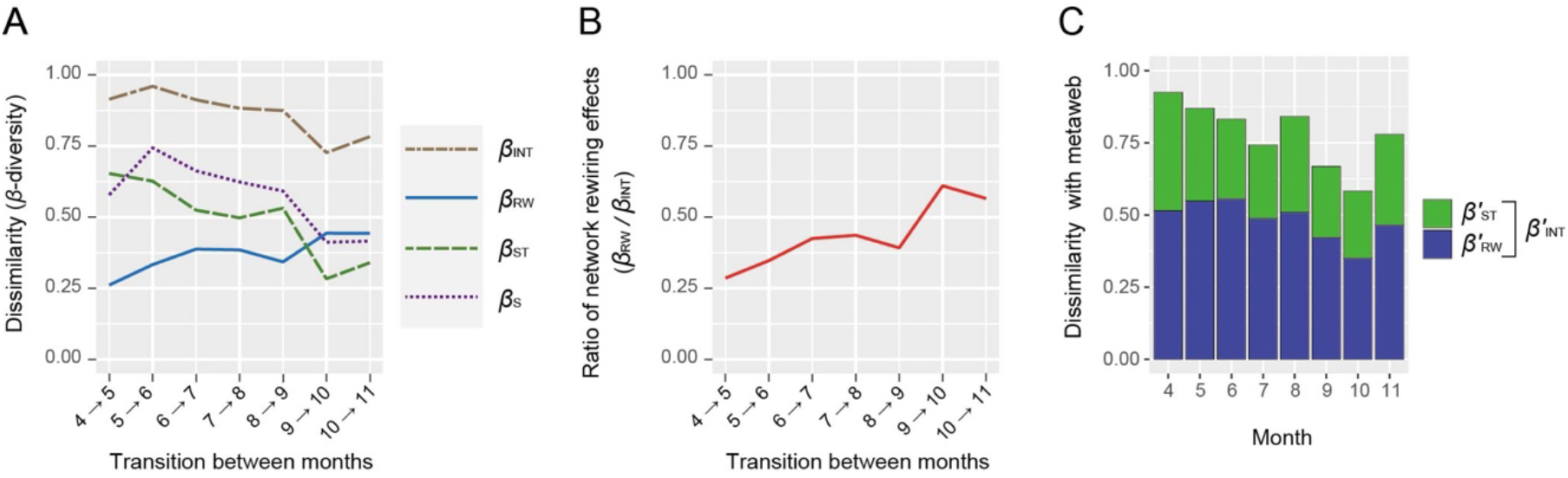
Network dissimilarity scores. (A) Changes in network architecture through time. Dissimilarity in network architecture between each pair of consecutive months (e.g., from April to May) are shown for each *β*-diversity component. (B) Ratio of interaction rewiring effects to total dissimilarity in network architecture. For each pair of consecutive months, relative contributions of interaction rewiring are evaluated by *β*_RW_/*β*_INT_. (C) Dissimilarity with the meta-network. For each pair of a realization (month) and the meta-network, dissimilarity in network architecture is shown. Note that dissimilarity in network architecture (*β*_INT_) consists of dissimilarity due to interaction rewiring (*β*_RW_) and that due to species turnover (*β*_ST_): i.e., *β*_INT_ = *β*_RW_ + *β*_ST_.

### Dissimilarity between each realization and the meta-network

We also found that network architecture of each month deviated considerably from that of the meta-network (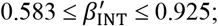; Fig. 3C). In all the months, effects of interaction rewiring 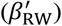 exceeded those of species turnover 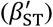 (Fig. 3C).

### Species contributions to network flexibility

Potential contributions to network flexibility 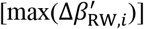 differed considerably among the spider species examined (Fig. 4A; Fig. S2). The presence of *Oxyopes sertatus* (Oxyopidae), which showed the highest contributions to network flexibility, was expected to increase the *β*-diversity component due to interaction rewiring 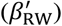 by 0.0587 at the maximum (Fig. 4A). Likewise, *Argiope bruennichi* (Araneidae) was inferred to increase 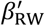by 0.0387 (Fig. 4A). In addition to those species, Salticidae sp.1, *Tetragnatha caudicula* (Tetragnathidae), *Pachygnatha tenera* (Tetragnathidae), *Neoscona adianta* (Araneidae), *Ebrechtella tricuspidata* (Thomisidae), and *Xysticus* sp. (Thomisidae) displayed relatively high contributions to network flexibility. We also found that *O. sertatus* and *A. bruennichi* had the greatest contributions to total network dissimilarity between realizations and the meta-network [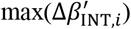; Fig. S3].

**Figure 4.**
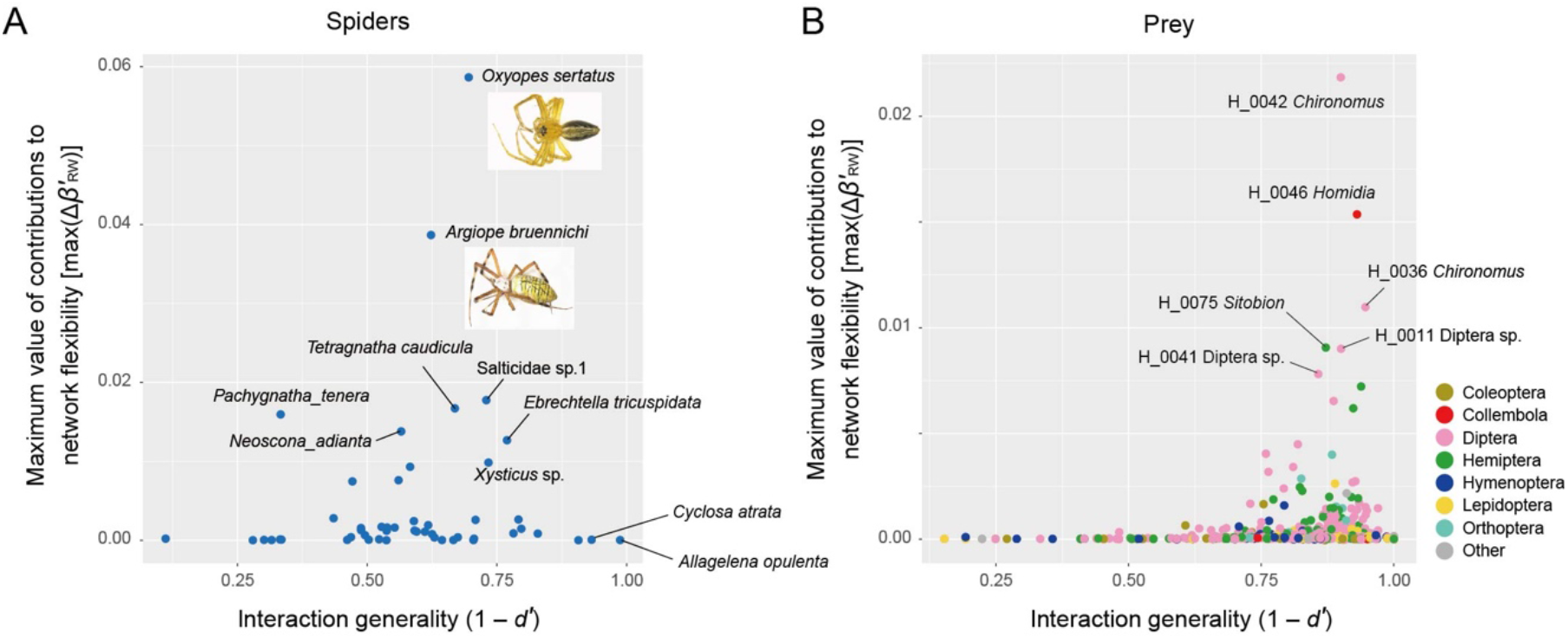
Contributions of each species to network flexibility. (A) Contributions of spider species to network rewiring. The maximum value of contributions to network rewiring effects 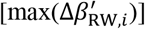 is shown for each spider species on the vertical axis. Species with higher 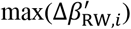 values have higher potential impacts on the flexibility of interaction networks. The horizontal axis indicates the interaction generality index (1 – *d*^′^, where *d*^′^ is standardized Kullback-Leibler divergence representing interaction specificity) calculated based on the meta-network including all the vertices (spider species and prey OTUs) and edges (interactions) observed across the eight months. (B) Contributions of prey OTUs to network rewiring.

We then found that most generalist predators did not necessarily have highest contributions to network flexibility. Specifically, when “interaction generality” to prey is evaluated based on the standardized Kullback-Leibler divergence of the meta-network matrix (33, 36), the two species with the highest contributions to network flexibility (*O. sertatus* and *A. bruennichi*) displayed moderate levels of interaction generality (0.696 and 0.623; Fig. 4A). This result indicates that species’ contributions through network dynamics can be independent of species properties along the axis of generalist–specialist continuums.

Among the 974 prey species/strains, those belonging to the genera *Chironomus* (Chironomidae; Diptera) and *Homidia* (Entomobrydae; Collembola) had the highest potential contributions to network flexibility (Fig. 4B; Fig. S2). These species also showed highest levels of interaction generality (0.900– 0.946). Thus, the prey species non-selectively consumed by a wide range of spider species could increase the flexibility of network architecture in the grassland ecosystem.

## Discussion

We here quantitatively evaluated the extent to which predator–prey interaction networks vary through time in the wild. Although there have been classic studies reporting entangled webs of consumer–victim interactions (37–39), our study, as far as we know, is the first to evaluate architectural flexibility of networks involving hundreds of species. As shown in this study, network architecture of predator–prey systems shift within short time windows due to seasonal turnover of species compositions (as represented by *β*_ST_) as well as due to rewiring of interactions (*β*_RW_). Without considering such flexibility of ecological networks, mechanisms determining community stability will never be fully understood. Thus, fueling feedback between empirical and theoretical studies beyond static view of interaction networks is the starting point for reorganizing our recognition of species coexistence mechanisms.

Comparing the magnitude of network architectural shifts between different types of species interactions is of particular importance for comprehensively understanding consequences of network dynamics. A pioneering work on plant–pollinator interactions (24) reported that total changes in network architecture could be large (*β*_INT_ > 0.50) across seasonal transitions and that changes due to interaction rewiring could consistently exceed those due to species turnover (*β*_RW_ > *β*_ST_). In our analysis on predator– prey interactions, total changes in network architecture were large as well (*β*_INT_ > 0.50), while balance between interaction rewiring and species turnover effects shifted across the seasons (Fig. 3A-B). These findings illuminate the working hypothesis that flexibility of network architecture is a basic property common to mutualistic and antagonistic interactions despite potential difference in factors driving such network architectural dynamics. Further accumulation of empirical studies is essential for systematically evaluating the commonality and diversity of species-rich network dynamics.

Based on the time-series analysis of predator–prey interactions, we developed a framework for evaluating species’ contributions to the flexibility of ecological network architecture. Theoretical studies have predicted that predators’ adaptive food choice (30), which is represented by dynamically changing interaction coefficients in mathematical models, can aid long-term stability of complex communities (i.e., communities with high species richness and connectance) (12). Without such adaptive foraging, community complexity is negatively associated with stability as suggested by May’s classic model (6), while with predators’ adaptive choice, complexity can promote community persistence (12). Consequently, species increasing network architectural flexibility are possibly keys to understand the reason why classic models do not explain empirical observations on positive relationship between community complexity and stability (40).

In our analysis on the spider–prey interactions, a sit-and-wait-type spider *O. sertatus* and a web-weaving spider *A. bruennichi* had much greater impacts on network flexibility than other species (Fig. 4A). Although spiders are often considered as generalist predators (41), having broad potential prey ranges does not necessarily guarantee high contributions of the species to network architectural plasticity. In other words, responsiveness to biotic/abiotic environmental changes is another essential factor to be explored through community dynamics.

Investigations of such “network coordinators” add a new dimension to the discussion of “keystoneness” in ecosystems (42). Since Paine’s seminal work (3, 43), keystone species, which impose great impacts on ecosystem-level dynamics (42), have been detected based on experimental exclusion of candidate species (28). However, pinpointing candidates of keystone species out of hundreds or thousands of species within ecosystems *per se* is basically an exhausting task. In this respect, network information provides bird’s-eye views for exploring potential keystone species (44–46), which should be subjected to detailed experimental investigations. The framework proposed in this study is expected to highlight predators imposing flexible top-down control on diverse prey and thereby promoting species coexistence at the lower trophic level (27, 28). It is also expected to illuminate prey species buffering biotic/abiotic perturbations by being flexibly consumed by diverse predator species. In advancing the application of this network-based approach, benchmark analyses quantifying the extent to which classic examples of keystone species [e.g., sea stars in intertidal communities (3, 43)] contribute to network flexibility is awaited.

The present constraint limiting our knowledge of network architectural dynamics is the scarcity of empirical datasets covering multiple time points (25). Therefore, it is worthwhile to examine whether some network indices deriving from static network analyses can be used as proxies for the network coordinator index. In this respect, network centrality metrics, such as degree and betweenness centralities (44, 45, 47), may provide a broadly applicable platform. However, we found that species with high network centralities do not necessarily have high potential contributions to network flexibility (Figs. S4-5). This result suggests that time-series analyses shed new lights on the organization of ecological networks and that more empirical datasets covering multiple time points are required to deepen our understanding of keystone species.

Further conceptual advances are necessary for systematically exploring species controlling ecological community dynamics and stability. Keystone species have been conventionally defined as species having disproportionately large impacts on ecosystems relative to its abundance (42). In this respect, it may be important to compare network coordinator scores among species with the same levels of abundance within a community dataset (Fig. S6). Furthermore, the assumption that keystone species can be replaced through time (33) would be an alternative basis for interpreting real community dynamics.

The guild or functional group of highlighted network coordinators is another important target of discussion. For example, the fact that possibly detritivores prey such as nonbiting midges (*Chironomus*) and springtails (*Homidia*) showed high contributions to network flexibility is of particular interest. This result leads to the hypothesis that stability of above-ground food webs is maintained by subsidy from below-ground ecosystems (48, 49). Thus, ecological roles of the potential network coordinators at the interface of different energy channels deserve future extensive studies (49, 50).

In the era of worldwide biodiversity loss and ecosystem degradation, highlighting keystone species is an essential task in conservation biology (26, 27, 46, 51). Considering concurrently escalating issues such as global warming, frequent extreme weather events, and environmental pollution, prioritized conservation efforts need to be directed to species buffering biotic/abiotic environmental changes within flexible webs of interactions. Given that high-throughput analyses of species interactions are becoming possible based on DNA metabarcoding (45), time-series analyses of interaction networks (52) will be widely applied as essential platforms in ecosystem conservation and restoration programs. Bird’s-eye views for exploring network coordinator species will advance both basic and applied sides of ecosystem sciences.

## Materials and Methods

### Dataset

We compiled the DNA metabarcoding dataset of spiders’ prey contents in a warm-temperate grassland located at Center for Ecological Research, Kyoto University, Japan (34°58’16.7”N 135; 57’32.3”E) (33). At the study site, spiders were haphazardly collected by sweeping with an insect net (diameter = 50 cm) on 3–5 days in the middle of each month from April to November, 2018 (168–441 spider individuals per month). Note that spiders were hard to sample in winter (from December to March) due to the inactivity of them as well as the lack of grasses to sweep. Because all the spider individuals (larger than 2 mm in body length) caught in the sweeping net were sampled, the collected specimens as a whole represented the species compositions of spiders at the study site in each month. In total, 2,224 spider samples representing 63 species were sampled across the eight months (33). Each spider sample was washed sequentially with distilled water, 70 % ethanol, and 100 % ethanol. Prey repertoires of each spider sample was then reconstructed based on the DNA metabarcoding (illumina amplicon sequencing) targeting the mitochondrial 16S rRNA region of Hexapoda (i.e., insects, springtails, etc.).

Prey DNA was detected from 1,556 out of the 2,224 spider samples examined (33). In the metabarcoding data, the presence (1) and absence (0) of each prey OTU in each spider sample (individual) was designated for each month. The information of each month was used to obtain a “species-level” matrix, in which cell entries represented the number of spider samples from which respective spider-Hexapoda OTU combinations were observed (i.e., prey detection counts). Across the eight months (realizations), 974 prey OTUs (defined with 97 % threshold identity of mitochondrial 16S rRNA sequences) belonging to 120 families were present in the dataset. The total number of spider–Hexapoda links was 2,247, representing 5,190 prey detection counts (33). The network of each month (realization) and the “meta-network” containing all the interactions observed across the eight months were visualized using the bipartite v.2.6-2 package of R 4.1.2.

### Network dissimilarity between realizations

For a pair of ecological communities (realizations), difference in network architecture could be expressed as *β*-diversity of network edge (link) compositions (25). The *β*-diversity index representing dissimilarity in interaction network architecture (*β*_INT_) could have two additive components, namely, dissimilarity in network architecture among species that occurred in both months (i.e., dissimilarity due to network rewiring; *β*_RW_) and dissimilarity in network architecture due to difference in species compositions between the months (i.e., dissimilarity due to species turnover; *β*_ST_) (24, 25). Therefore, the relationship between *β*_INT_, *β*_RW_, and *β*_ST_ can be expressed as:

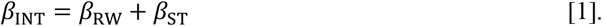

Dissimilarity in network architecture can be then calculated based on various types of *β*-diversity metrics. When Sørensen’s metric for binary data formats, for example, is used to evaluate total dissimilarity between networks (realizations) **M** and **N**, *β*_INT_ can be expressed as

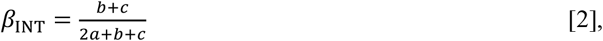

where *a* denotes the number of items (network edges) commonly observed in the two networks (**M** and **N**) compared, *b* is the number of items unique to network **M**, and *c* is the number of items unique to network **N** (25). Likewise, dissimilarity in network architecture due to rewiring is calculated by redefining networks to be compared. Specifically, by focusing on network components that contain only species shared between the two networks (realizations), subset networks **M**_shared_ and **N**_shared_ are obtained. *β*_RW_ is then calculated as

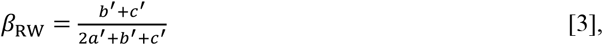

where *a*^′^ denotes the number of items (network edges) commonly observed in the two networks (**M**_shared_ and **N**_shared_) compared, *b*^′^ is the number of items unique to network **M**_shared_, and *c*^′^ is the number of items unique to network **N**_shared_. In this framework proposed by Poisot *et al*. (25) (hereafter, framework 1), *β*_ST_ is then calculated by substracting *β*_RW_ from *β*_INT_: i.e.,

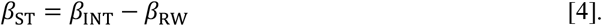

This indirect estimation of *β*_ST_ is not guaranteed if the additivity of the *β*-diversity components is not met (34). Therefore, for accurately partitioning network architectural dissimilarity into effects of network rewiring and those of species turnover, an alternative framework of *β*-diversity calculation has been proposed (34, 35) (hereafter, framework 2). Among the studies, Fründ (34) has proposed to calculate both *β*_RW_ and *β*_ST_ directly with Sørensen’s *β*-diversity measure as follows:

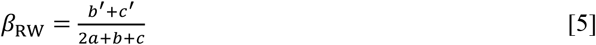

and

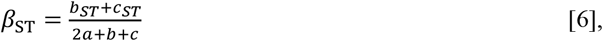

where *b*_*ST*_ and *c*_*ST*_ are obtained as *b* − *b*^′^ and *c* − *c*^′^, respectively. Because the two indices are calculated with the common denominator (i.e., 2*a* + *b* + *c*), their sum is always equal to *β*_*WN*_ as follows:

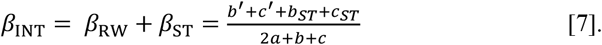

In this study, we used framework 2 for partitioning network rewiring and species turnover effects in calculating dissimilarity in interaction network architecture. Meanwhile, we performed a comparative analysis based on framework 1 as shown in Figures S7-8. In both frameworks, prey detection counts in the input data were converted into proportions using the “proportions=TRUE” option in the “betalinkr” function (34) of the R bipartite package (53). Each dissimilarity indices were then calculated based on Bray-Curtis metric of *β*-diversity using the “betalinkr” function.

### Transitions of network architecture

Dissimilarity in network architecture through the time-series was evaluated based on the abovementioned *β*-diversity indices. For each pair of consecutive months (e.g., from April to May), total dissimilarity in interaction network architecture (*β*_INT_), dissimilarity in network architecture due to rewiring (*β*_RW_), and dissimilarity in network architecture due to species turnover (*β*_ST_) were evaluated. In addition to dissimilarity in network architecture, dissimilarity in species compositions (*β*_S_) were calculated based on Bray-Curtis *β*-diversity.

### Dissimilarity between each realization and the meta-network

We also calculated dissimilarity in network architecture between each realization (each month’s network) and the meta-network representing all the spider–prey interactions detected across the eight months (Fig. 1). Specifically, total dissimilarity in network architecture, dissimilarity in network architecture due to rewiring, and dissimilarity in network architecture due to species turnover were calculated as 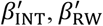, and 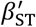, respectively, for each pair of a realization and the meta-network (Fig. 1). Likewise, dissimilarity in species compositions between a realization and the meta-network 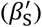 was calculated.

### Species contributions to network flexibility

To evaluate potential contributions of each species to flexibility of network architecture, we examined the extent to which dissimilarity in network architecture between a realization and the meta-network could change due to interaction rewiring effects of a target species. We then developed 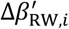 index, which was defined as follows:

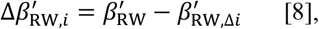

where 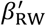was the original value of dissimilarity in network architecture due to rewiring as defined above, and 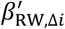denoted the simulated value of dissimilarity calculated by removing species *i* from the dataset. By definition, this 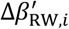index can be calculated for each species in the dataset in each pair of a realization (month) and the meta-network. Therefore, for each species *i*, we used the maximum value of 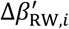across the time-series [i.e., 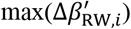] as a measure of potential magnitude of contributions to network architectural flexibility: a positive value of 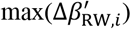 indicated positive contributions of a target species to network flexibility.

For further evaluating roles of each species to total dissimilarity in network architecture and dissimilarity through species turnover, we also calculated contributions of species *i* to 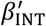 and 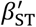as defined below:

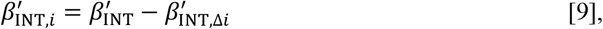

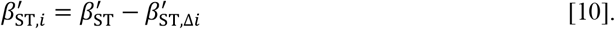

Likewise, contributions of species *i* to community compositional dissimilarity 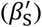 was calculated as follows:

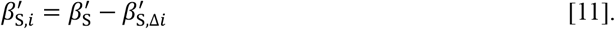

Maximum values of those indices across the time-series were calculated as well to evaluate potential contributions of each species.

### Interaction generality

To evaluate basic specificity/generality of spiders to their prey, we examined the extent to which each spider species interacted specifically to its prey based on the *d*^′^ index of standardized Kullback-Leibler divergence (36). As *d*^′^ score varies from 0 (minimum specificity given a species abundance within the community) to 1 (maximum specificity given a species abundance), the value 1 – *d*^′^ can be used as a measure of the extent to which a spider species interacted with broad ranges of prey (“interaction generality”) (33). The interaction generality index was calculated, respectively, for the data matrix of each month and that of the meta-network. The interaction generality of each prey OTU to predator (spider) species was calculated as well.

### Network centrality

To evaluate the extent to which each spider species or prey OTUs was located at the core position within a network, we calculated degree and betweenness centrality. Degree centrality was defined as the number of edges connected to the target vertex (species or OTU). The obtained degree centrality was then normalized by dividing it by *N* – 1, where N is the number of vertices within the target network. Betweenness centrality is a measure of the degree to which a given vertex is located within the shortest paths connecting pairs of other vertices in a network (47). Scores of betweenness were normalized for within each network so that they varied from 0 (occupation at marginal positions within a network) to 1 (occupation at shortest paths for all pairs of vertices) using the igraph v.1.3.0 package (54) of R.

## Code availability

All the codes used for the analyses are available from the GitHub repository (https://github.com/hiro-toju/network_flexibility_spiders) [To be made publicly available after the acceptance of the paper].

## Acknowledgments

This work was financially supported by JSPS Grant-in-Aid for Scientific Research (18H04009) and JST FOREST (JPMJFR2048) to H.T..

## Notes

### Competing Interest Statement

The authors have declared no competing interest.

