## Supplementary Figures for "Dynamics of interaction networks and species’ contributions to community-scale flexibility"

*Hirokazu Toju

**This PDF file includes:**

Figures S1 to S8

### Spider species

| | |
|---|---|
| Age_sil | *Agelena silvatica* |
| All_opu | *Allagelena opulenta* |
| Ara_eju | *Araneus ejusmodi* |
| Ara_sp. | *Araniella* sp. |
| Ara_sp.1 | *Araneidae* sp.1 |
| Ara_sp.2 | *Araneidae* sp.2 |
| Arg_amo | *Argiope amoena* |
| Arg_bon | *Argyrodes bonadea* |
| Arg_bru | *Argiope bruennichi* |
| Ari_cyl | *Ariamnes cylindrogaster* |
| Bat_sp. | *Bathyphantes* sp. |
| Che_jap | *Cheiracanthium japonicum* |
| Clu_sp. | *Clubiona* sp. |
| Cyc_atr | *Cyclosa atrata* |
| Cyc_sed | *Cyclosa sedeculata* |
| Cyr_nag | *Cyrtarachne nagasakiensis* |
| Dia_sub | *Diaea subdola* |
| Dol_sp. | *Dolomedes* sp. |
| Ebr_tri | *Ebrechtella tricuspidata* |
| Epi_sp. | *Episinus* sp. |
| Gib_abs | *Gibbaranea abscissus* |
| Hyp_pyg | *Hypsosinga pygmaea* |
| Lar_arg | *Larinia argiopiformis* |
| Leu_bla | *Leucauge blanda* |
| Leu_cel | *Leucauge celebesiana* |
| Lin_sp.1 | *Linyphiidae sp.1* |
| Lin_sp.2 | *Linyphiidae sp.2* |
| Lin_sp.3 | *Linyphiidae sp.3* |
| Lin_sp.4 | *Linyphiidae sp.4* |
| Lin_sp.5 | *Linyphiidae sp.5* |
| Lyc_sp.1 | *Lycosidae sp.1* |
| Lyc_sp.2 | *Lycosidae sp.2* |
| Men_elo | *Mendoza elongata* |
| Mim_sp. | *Mimetus* sp. |
| Myr_sp. | *Myrmarachne* sp. |
| Neo_adi | *Neoscona adianta* |
| Neo_mel | *Neoscona mellotteei* |
| Neo_nau | *Neoscona nautica* |
| Neo_scy | *Neoscona scylloides* |
| Ner_rad | *Neriene radiata* |
| Oxy_bad | *Oxyopes badius* |
| Oxy_ser | *Oxyopes sertatus* |
| Oxy_str | *Oxytate striatipes* |
| Pac_qua | *Pachygnatha quadrimaculata* |
| Pac_ten | *Pachygnatha tenera* |
| Par_jap | *Parasteatoda japonica* |
| Par_sp. | *Pardosa* sp. |
| Phi_sp. | *Philodromus* sp. |
| Sal_sp.1 | *Salticidae* sp.1 |
| Sal_sp.2 | *Salticidae* sp.2 |
| Sal_sp.3 | *Salticidae* sp.3 |
| Sal_sp.4 | *Salticidae* sp.4 |
| Sal_sp.5 | *Salticidae* sp.5 |
| Sal_sp.6 | *Salticidae* sp.6 |
| Sit_sp. | *Sitticus* sp. |
| Tet_cau | *Tetragnatha caudicula* |
| Tet_pra | *Tetragnatha praedonia* |
| Tet_squ | *Tetragnatha squamata* |
| The_sp. | *Theridiidae* sp. |
| Tho_lab | *Thomisus labefactus* |
| Tri_cla | *Trichonephila clavata* |
| Xys_sp. | *Xysticus* sp. |

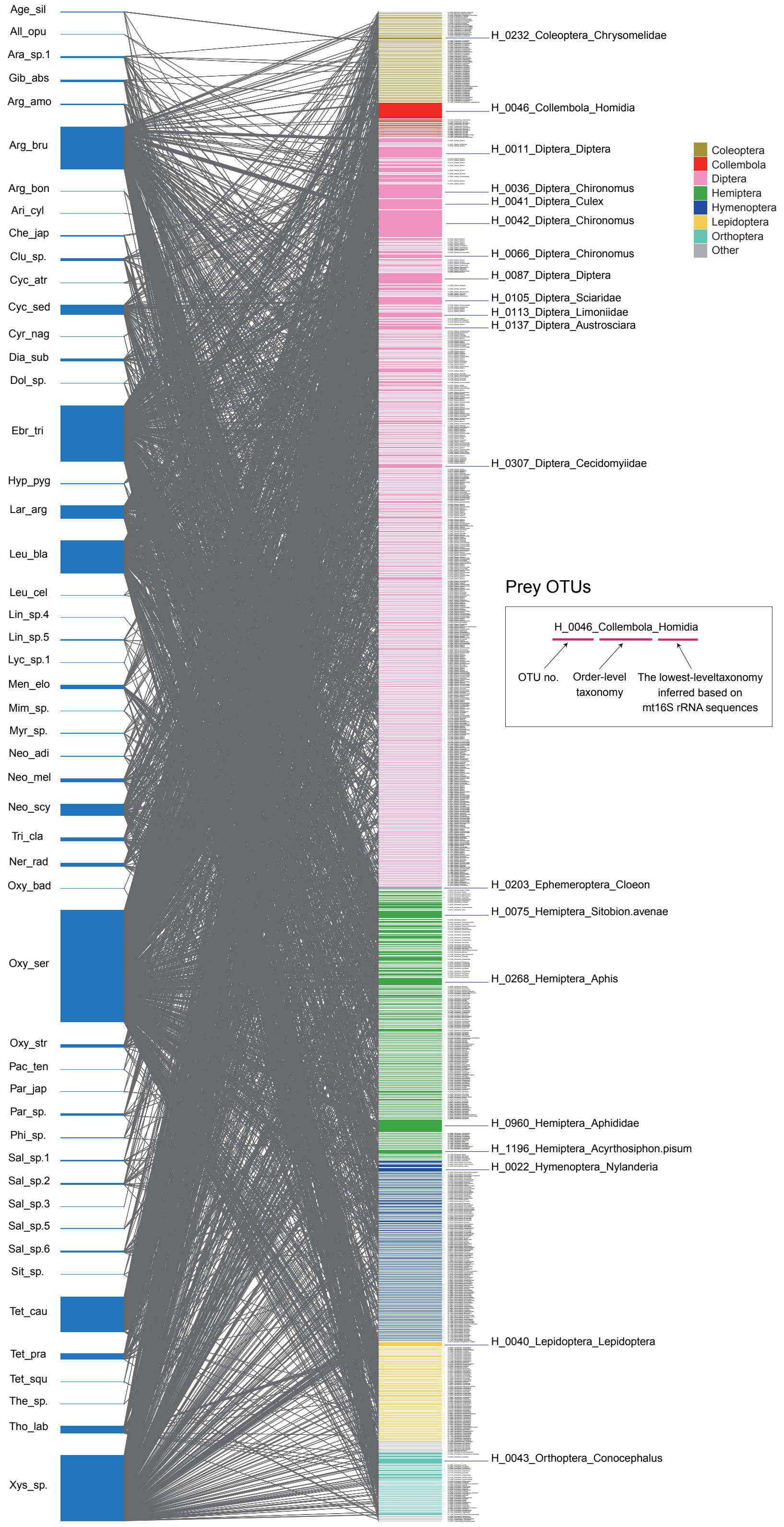


**Fig. S1.** Details of the spider–prey meta-networks. All the spider–prey interactions observed from April to November are included in the meta-network. Spider species and prey Hexapoda OTUs are shown in the left and right, respectively.

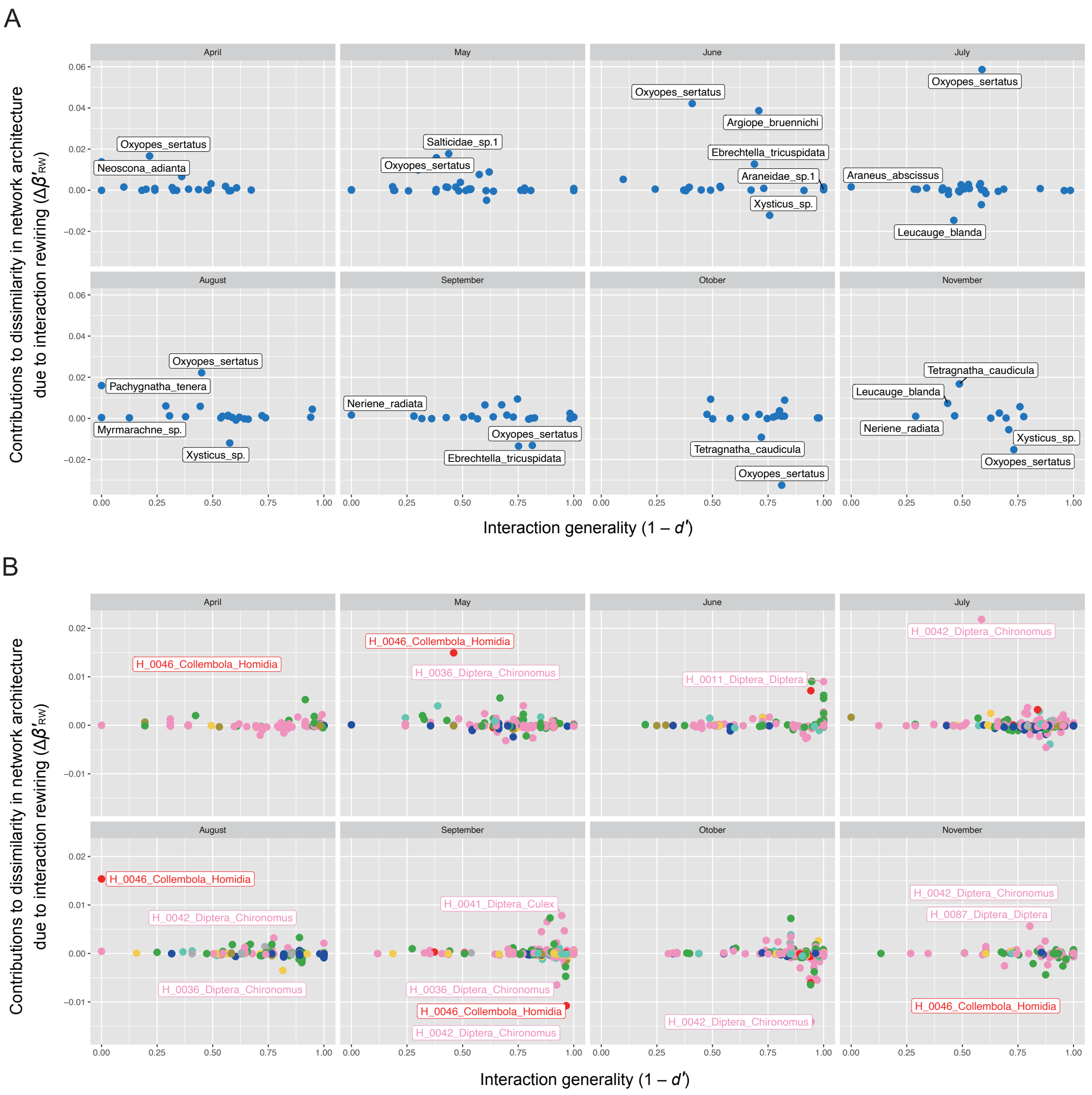


**Fig. S2.** Contributions of each species to dissimilarity in network architecture due to interaction rewiring. For each month, dissimilarity in network architecture due to interaction rewiring was calculated against the meta-network (Fig. 2A). Contributions of each spider species or prey OTUs to the dissimilarity levels ($\Delta\boldsymbol{\beta}'_{\mathbf{RW},i}$) were then evaluated as illustrated in Fig. 1B. (A) Spider species. (B) Prey OTUs.

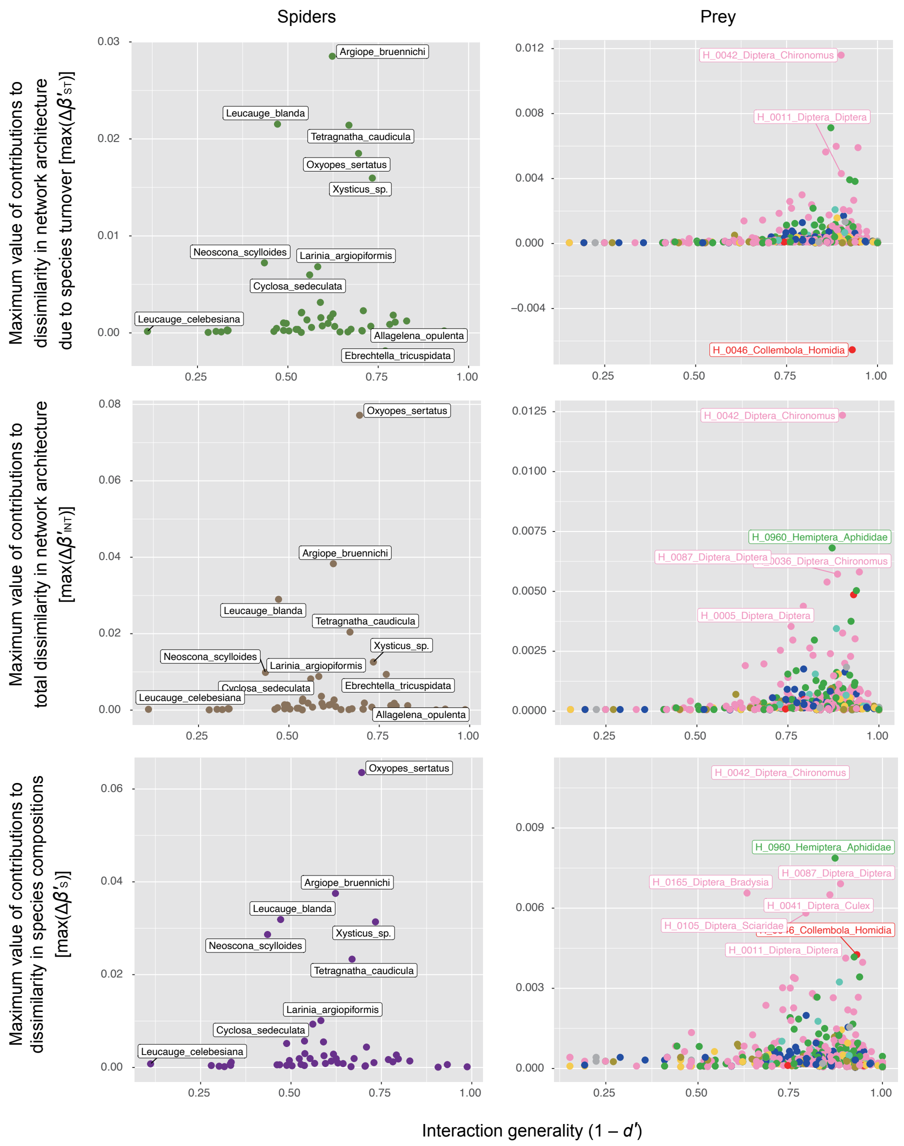


**Fig. S3.** Contributions of each species to various dissimilarity indices. In line with the maximum values of contributions to network rewiring effects [$\max(\Delta\boldsymbol{\beta}'_{\mathbf{RW},i})$] shown in Figure 4, the maximum values of contributions to dissimilarity in network architecture due to species turnover [$\max(\Delta\boldsymbol{\beta}'_{\mathbf{ST},i})$], the maximum values of contributions to total dissimilarity in network architecture [$\max(\Delta\boldsymbol{\beta}'_{\mathbf{INT},i})$], and the maximum values of contributions to dissimilarity in species compositions [$\max(\Delta\boldsymbol{\beta}'_{\mathbf{S},i})$] are shown respectively for spiders and prey.

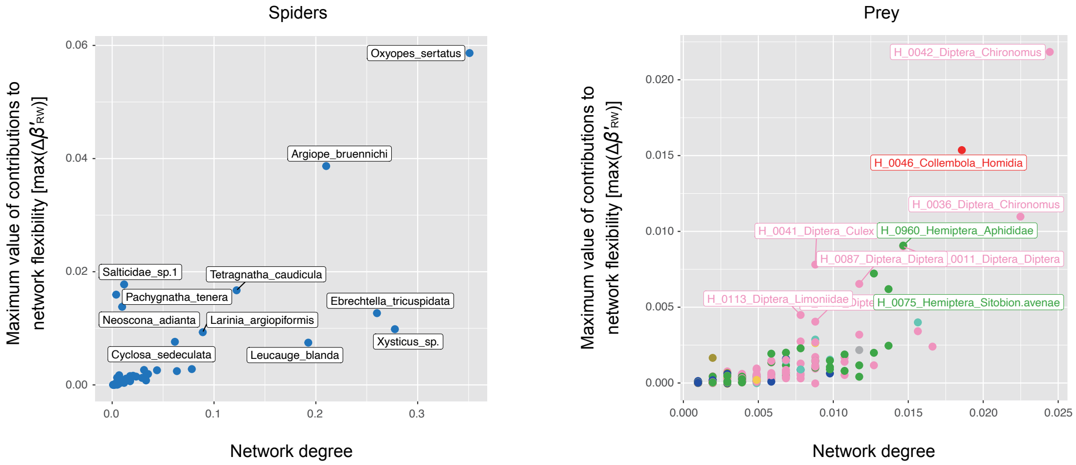


**Fig. S4.** Degree centrality and contributions to network flexibility. The maximum values of contributions to network rewiring effects $[\max(\Delta\boldsymbol{\beta}'_{\mathbf{RW},\boldsymbol{i}})]$ (Fig. 4) are plotted against the axis of degree centrality within the meta-network (Fig. 2A).

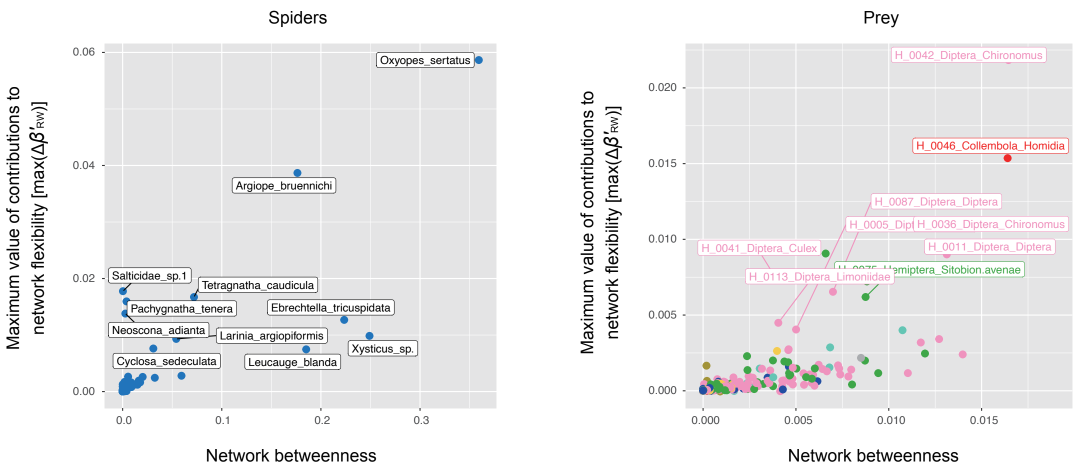


**Fig. S5.** Betweenness centrality and contributions to network flexibility. The maximum values of contributions to network rewiring effects [max($\Delta\boldsymbol{\beta}'_{\mathbf{RW},i}$)] (Fig. 4) are plotted against the axis of betweenness centrality within the meta-network (Fig. 2A).

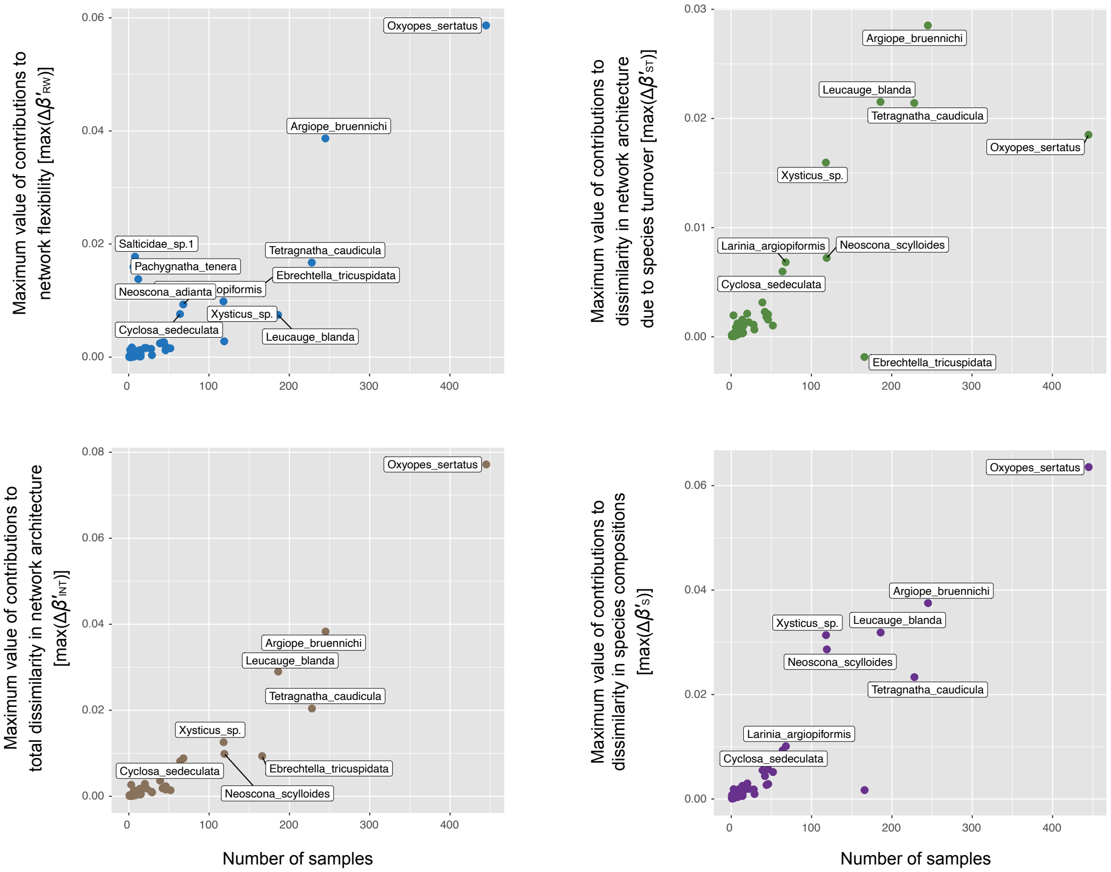


**Fig. S6.** Number of samples and contribution indices. The maximum values of contributions to network rewiring effects [max($\Delta\boldsymbol{\beta}'_{\mathbf{RW},i}$)], the maximum values of contributions to dissimilarity in network architecture due to species turnover [max($\Delta\boldsymbol{\beta}'_{\mathbf{ST},i}$)], the maximum values of contributions to total dissimilarity in network architecture [max($\Delta\boldsymbol{\beta}'_{\mathbf{INT},i}$)], and the maximum values of contributions to dissimilarity in species compositions [max($\Delta\boldsymbol{\beta}'_{\mathbf{S},i}$)] are plotted against the axis indicating the number of collected spider samples.

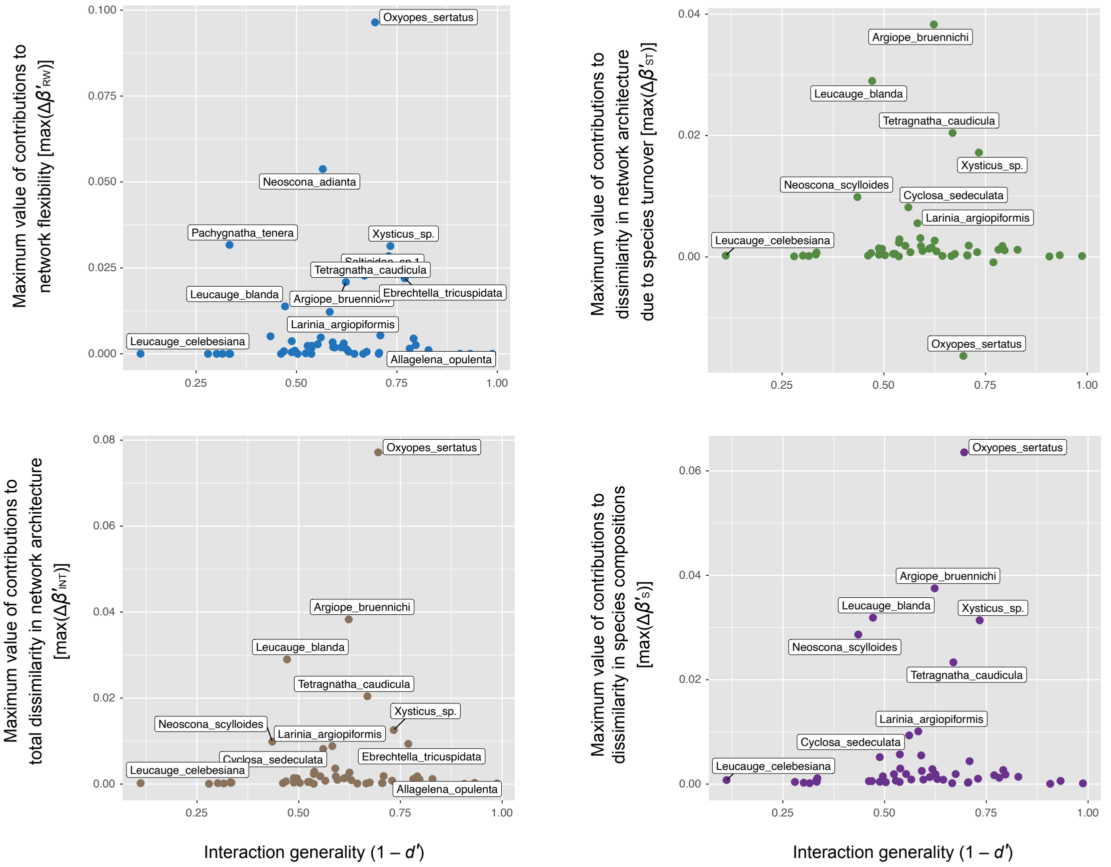


**Fig. S7.** Contribution indices calculated based on framework 1 (spiders). In the main body of the paper and Figures 2-6, the $\beta$-diversity-based indices were calculated based on framework 2, in which common denominators were used in the calculation of $\Delta\boldsymbol{\beta}_{\mathbf{RW}}$ and $\Delta\boldsymbol{\beta}_{\mathbf{ST}}$. For comparison with framework 2, the contribution indices were re-calculated based on framework 1, in which $\boldsymbol{\beta}_{\mathbf{ST}}$ was defined as $\boldsymbol{\beta}_{\mathbf{INT}} - \boldsymbol{\beta}_{\mathbf{RW}}$.

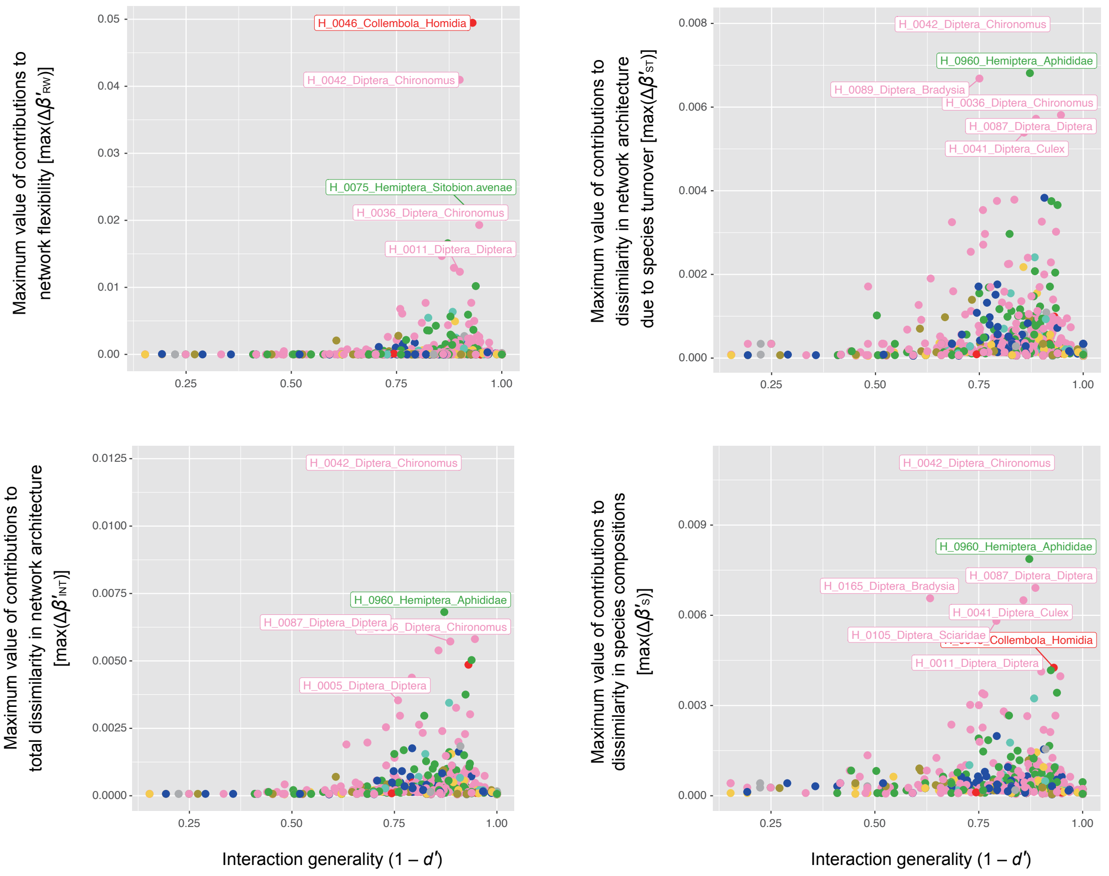


**Fig. S8.** Contribution indices calculated based on framework 1 (prey). In the main body of the paper and Figures 2-6, the $\beta$-diversity-based indices were calculated based on framework 2, in which common denominators were used in the calculation of $\Delta\boldsymbol{\beta}_{\mathbf{RW}}$ and $\Delta\boldsymbol{\beta}_{\mathbf{ST}}$. For comparison with framework 2, the contribution indices were re-calculated based on framework 1, in which $\boldsymbol{\beta}_{\mathbf{ST}}$ was defined as $\boldsymbol{\beta}_{\mathbf{INT}} - \boldsymbol{\beta}_{\mathbf{RW}}$.
